# Synchronized photoactivation of T4K rhodopsin causes a chromophore-dependent retinal degeneration that is moderated by interaction with phototransduction cascade components

**DOI:** 10.1101/2024.02.27.582380

**Authors:** Beatrice M. Tam, Paloma Burns, Colette N. Chiu, Orson L. Moritz

## Abstract

Multiple mutations in the *Rhodopsin* gene cause sector retinitis pigmentosa in humans and a corresponding light-exacerbated retinal degeneration (RD) in animal models. Previously we have shown that the *rhodopsin* mutation T4K requires photoactivation to exert its toxic effect. Here we further investigated the mechanisms involved in rod cell death caused by T4K rhodopsin in *Xenopus laevis*. In this model, RD was prevented by rearing animals in constant darkness but surprisingly also in constant light. RD was maximized by light cycles containing at least one hour of darkness and 20 minutes of light exposure, light of intensity 750 lux or greater, and by sudden light onset. Under conditions of frequent light cycling, RD occured rapidly and synchronously, with massive shedding of ROS fragments into the RPE initiated within hours, and subsequent death and phagocytosis of rod cell bodies. RD was minimized by reduced light levels, pre-treatment with constant light, and gradual light onset. RD was prevented by genetic ablation of the retinal isomerohydrolase RPE65, and exacerbated by ablation of phototransduction components GNAT1, SAG, and GRK1. Our results indicate that photoactivated T4K rhodopsin is toxic, that cell death requires synchronized photoactivation of T4K rhodopsin, and that toxicity is mitigated by interaction with other rod outer segment proteins regardless of whether they participate in activation or shutoff of phototransduction. In contrast, RD caused by P23H rhodopsin does not require photoactivation of the mutant protein, as it was exacerbated by RPE65 ablation, suggesting that these phenotypically similar disorders may benefit from different treatment strategies.

**Significance:** A large number of *rhodopsin* mutations are linked to the inherited degenerative disease retinitis pigmentosa. Although the end result in each case is the loss of photoreceptor cells and blindness, not all of these mutations cause cell death via the same mechanism. In order to design and test treatment therapies that target the disease at points as upstream as possible in the process, we require detailed understanding of the range and nature of these disease mechanisms. This study using a transgenic *Xenopus laevis* model has extended our understanding of how T4K rhodopsin and related mutations cause rod cell photoreceptor death via a phototoxic product, and how this mechanism differs from the more extensively researched protein misfolding mechanism underlying cell death caused by P23H rhodopsin.

## Introduction

Retinitis pigmentosa (RP) is a genetically inherited retinal degeneration (RD) with many different inheritance patterns, including dominant, X-linked, and recessive (Hartong et al., 2006). Autosomal dominant (ad) RP is often caused by mutations in the *Rhodopsin* (*RHO*) gene (Daiger et al., 2013). Cideciyan and others have described phenotypically distinct forms of RP caused by *RHO* mutations; particularly intriguing is “type B1” or “sector RP”, in which the retina is non-uniformly affected, suggesting both the possibility of an environmental influence, and the potential for preventative treatment (Li et al., 1994; Cideciyan et al., 1998). Generally, the lower retina (which receives greater illumination from overhead light sources) is more severely affected, suggesting that the environmental influence is light.

A large body of research has focused on the study of light-induced retinal degeneration in wild type rodent models, with the goal of uncovering downstream pathogenic mechanisms in common with those involved in hereditary retinal degenerations (Organisciak and Vaughan, 2010; Grimm and Remé, 2013). A number of paradigms have been described but in general, the approaches can be summarized as exposing animals to unusually long or intense light exposures. (NOELL et al., 1966; LaVail et al., 1987; Vihtelic and Hyde, 2000; Marc et al., 2008; Grimm and Remé, 2019). These studies have led to the understanding that light-induced toxicity is directly mediated by rhodopsin and is modulated by the rate of rhodopsin regeneration, which itself can be modulated by polymorphisms in the retinoid isomerase, RPE65, as well as the wavelength/quality of light (Grimm et al., 2000, 2001; Wenzel et al., 2001, 2005). To complement this work, many animal models inherited RDs also exist, allowing mutation-specific pathogenic mechanisms to be studied.

Much of the research into adRP mechanisms has focused on the relatively prevalent *RHO* mutation P23H (Dryja et al., 1990), which is associated with sector RP. In cell culture this mutation promotes misfolding of rod opsin that is mitigated by binding of 11-cis retinal chromophore, which acts as a “pharmacological chaperone” to stabilize the protein (Noorwez et al., 2003; Athanasiou et al., 2018). Thus, depletion of 11-cis retinal by greater light exposure of the lower retina provides a plausible explanation for the sector RP phenotype. Similarly, light exposure promotes RD caused by P23H rhodopsin in transgenic *X. laevis* (Tam and Moritz, 2007), as well as transgenic *Drosophila*, swine, mice and rats (Organisciak et al., 2003; Galy et al., 2005; Scott et al., 2017; Orlans et al., 2019), while dietary deprivation of vitamin A, the chromophore precursor, promotes RD (Tam et al., 2010). Pharmacological chaperones have been proposed as therapeutic interventions for RP (Mendes and Cheetham, 2008; Chen et al., 2014; Behnen et al., 2018).

Another *RHO* mutation that causes sector RP in humans is T4K (van den Born et al., 1994), which disrupts one of two N-linked glycosylation sites in the N-terminus of the protein (Fukuda et al., 1979; Kaushal et al., 1994; Tam and Moritz, 2009). In an animal model the resulting RD is similarly mitigated by dark rearing (Tam and Moritz, 2009; Tam et al., 2014), and this is also the case for other mutations that disrupt N-linked glycosylation, including T17M and N15S (Tam and Moritz, 2009). However, the pathogenic mechanisms of T4K and P23H rhodopsins are very different. T4K does not cause a biosynthetic defect and thus the mutant protein is not retained in the ER. Moreover, RD can be prevented by a mutation in ‘cis’ that prevents chromophore binding and photoactivation as well as by vitamin A deprivation (Tam et al., 2014), whereas these interventions have the opposite effect of exacerbating RD in P23H animals (Tam and Moritz, 2007; Tam et al., 2010). Thus these two models represent distinct pathways for disease and therapeutic intervention.

In this study, we examine lighting paradigm details and genetic interactions that promote RD in *RHO* T4K transgenic *X. laevis*. We demonstrate that this RD has a novel requirement for cyclic light, and more rapid cycling promotes RD. We develop new methods for identifying genetic interactions in *X. laevis* to show that disruption of the *RPE65* gene prevents RD, and rhodopsin interaction partners from the phototransduction cascade are protective, regardless of whether they are involved in phototransduction activation or inactivation. We also show that unlike RD caused by *RHO* P23H, this RD involves minimal ultrastructural changes to rod outer and inner segments.

Our study indicates that distinct mechanisms underly different genetic causes of sector RP, which may therefore respond very differently to environmental influences or treatments. Recommendations can be made for *RHO* T4K patients specifically and clinical trial design generally. We also note similarities between RD caused by T4K rhodopsin and RD caused by intense or prolonged light exposure that suggest mechanistic relationships between these paradigms.

## Methods

### Animal husbandry

*X. laevis* tadpoles carrying a human T4K *RHO* transgene were generated by mating heterozygous transgenic animals with WT animals as in previous studies (Tam et al., 2014). Tadpoles were housed in clear plastic tanks at 18°C in 10 mM NaCl, 0.2 mM KCl, 0.1 mM MgCl2, 0.2 mM CaCl2. All experiments adhered to the ARVO statement for the Use of Animals in Ophthalmic and Vision Research.

### Light exposure

Unless otherwise stated, standard lighting conditions were 1700-2000 lux, 12 hours on, 12 hours off. Variant lighting conditions were achieved by placing the tanks in foil-lined cardboard boxes lit from above with white LED lights for which the timing and intensity could be varied. For each described experiment, all tadpoles were siblings from the same mating and approximately 50% of tadpoles were WT and 50% were transgenic. For any given lighting regimen, the experimental and control animals were housed together and subjected to identical conditions (lighting, temperature, feeding etc).

### Dot blot immunoassay

Normally developed tadpoles were sacrificed at the specified timepoints. One eye was solubilized in 100 μl of a 1:1 mixture of phosphate-buffered saline and SDS-PAGE loading buffer containing 1mM ethylenediaminetetraacetic acid and 100 μg/ml phenylmethylsulfonyl fluoride. Dot blot assays were performed on solubilized eyes as described by Tam et al. (Tam et al., 2006) and probed with anti-rod opsin antibodies monoclonal B630N (binds both *X. laevis* and human rod opsin) or monoclonal 1D4 (binds mammalian rod opsin only, used for genotyping) (1:10 dilution of tissue culture supernatant, and 1:5000 dilution of purified antibody respectively). Secondary antibody was IR-dye800-conjugated goat anti-mouse used at 1:10000 of 1 mg/ml solution (LI-COR Biosciences). A LI-COR Odyssey imaging system was used to image and quantify integrated intensity of blot signals. Log-transformed dot blot data was analyzed by 2-way ANOVA and Tukey post-hoc tests, using SPSS software (IBM).

### Western blot

Western blots were performed as previously described (Tam and Moritz, 2007). Briefly, solubilized eye samples prepared as described above for dot blot were separated on 10% SDS-PAGE gels, transferred overnight to immobilon P-membranes, and probed with the indicated antibodies and secondary antibodies as for dot blots. A LI-COR Odyssey imaging system was used to image the blots, and ImageJ software was used to quantify blot signals.

### Confocal microscopy

Samples were prepared for confocal microscopy as previously described (Tam et al., 2014). Briefly, contralateral eyes were fixed in 4% paraformaldehyde buffered with 0.1 M sodium phosphate pH 7.4. Fixed eyes were infiltrated with 20% sucrose for 16-48 hours, embedded in OCT (Sakura Finetek), frozen, and stored at −80°C before sectioning. 12 µm slices were cut using a cryostat and labelled overnight with primary antibody, followed by Cy3-conjugated anti-mouse and/or anti-rabbit secondary antibody (Jackson Research, West Grove, PA, USA), or Alexa647-conjugated anti-rabbit secondary antibody (for double-labeling protocols), and Alexa488-conjugated wheat germ agglutinin (WGA) and Hoechst 33342 counterstains. The following antibodies were used: monoclonal B630N (binds all rod opsin) monoclonal 2B2 (binds mammalian rod opsin), monoclonal 1D4 (binds mammalian rod opsin), polyclonal YZS437 (RPE65) (Wen et al., 2019), polyclonal anti-GNAT1 (a rabbit polyclonal anti-transducin antibody that was produced against the *X. laevis* GNAT1 protein expressed in and purified from *E. coli*), monoclonal Xarr1-6 anti-SAG (Peterson et al., 2003) and monoclonal G8 anti-GRK1 (Zhao et al., 1998)(Abcam). Images were obtained using a Zeiss 510 meta confocal microscope equipped with a 40X N.A. 1.2 water-immersion objective (Carl Zeiss, Oberkochen, Baden-Württemberg, Germany). Figures were assembled in Adobe Photoshop. Antibody labeling signal was adjusted linearly and equally across panels, and WGA and Hoechst signals were adjusted non-linearly to improve fine detail.

### Electron microscopy

Samples were prepared as previously described (Tam et al., 2015). Briefly, eyes were fixed in 4% paraformaldehyde+1% glutaraldehyde in 0.1 M phosphate buffer pH 7.4 at 4°C for ≥24 h. Fixed eyes were then infiltrated with 2.3 M sucrose in 0.1 M phosphate buffer, embedded in OCT (Sakura Finetek), cryosectioned, and 20 µm slices were collected on gelatin-coated slides. Optimally oriented sections were washed with 0.1 M sodium cacodylate and then stained for 30 min with 1% osmium tetroxide. After staining, sections were dehydrated in increasing concentrations of anhydrous ethanol and infiltrated with Eponate 12 resin (Ted Pella Inc., Redding, CA). Trimmed Beem® capsules were placed over the section on the slide, filled with resin and polymerized 16–24 h at 65°C. Blocks were separated from slides, and ultrathin sections (silver-gray; 50–70 nm) were cut with a diamond knife and collected on 0.5% Formvar-coated nickel slot grids. Sections were stained with saturated methanolic uranyl acetate (5 min) and Venable and Coggeshall’s lead citrate (0.25%, 5 min). Imaging was performed with a Hitachi 7600 TEM at 80 kV. In some cases, panels were assembled from multiple images using the “photomerge” function in Adobe photoshop. Tinting was manually added to some images using Adobe Photoshop.

### Gene editing

CRISPR-based gene editing was carried out essentially as previously described (Feehan et al., 2019). Eggs were obtained from heterozygous transgenic females and fertilized in-vitro using WT male sperm, generating a population of 50% transgenic embryos. Single-celled embryos were injected with Cas9 mRNA with or without the addition of sgRNA to generate populations of edited or non-edited embryos. An eGFP mRNA was co-injected, and used to identify successfully injected embryos at 24 hours post-fertilization by fluorescence microscopy. Cas9, eGFP and sgRNAs were generated and transcribed *in vitro* as previously described (Feehan et al., 2019) using the vectors pBluescript SK+ (Stratagene) and pDR274 (gift of Keith Joung). The target sequences of the sgRNAs were as follows: *RPE65:* 5’-GGGGAACTGAACAACAACCT-3’. *GNAT1*: 5’-ATGGAAAGACACGGGTATCC-3’(1) 5’-GGTATCATTGAGACACAGTT-3’(2); *SAG.L*: 5’-GGTTTAGGCCCTGGCTGATC-3’, *SAG.S*: 5’-GGCATTGACCGCACCGTCAT-3’; *GRK1*: 5’-GGATCGTTTGCGGAAAGCTA-3’; N (control sgRNA): 5’ACTTGCCTTGTCTCCCAGGG-3’. Note that procedures for gene editing of *RPE65* and *GNAT1* were identical to those described previously (Wen et al., 2019). Gene editing was confirmed by PCR of the edited locations, and Sanger sequencing of the PCR products, with procedures identical to those previously described (Feehan et al., 2017; Wen et al., 2019).

### Electroretinography

Electroretinography was performed as previously described (Vent-Schmidt et al., 2017). Briefly, animals aged 6-12 weeks were dark-adapted overnight, and further procedures were done under red light. Animals were anesthetized using 0.5% buffered Tricaine solution, and mounted on a gold ECG electrode. A chlorided silver wire electrode mounted in a glass capillary was placed on the cornea, and the electrodes were connected to the headstage of a model 1800 AC amplifier (AM systems). The output of the amplifier was connected to the input of an Espion electroretinography unit (Diagnosys). The Ganzfeld dome of the Espion unit was lowered over the tadpole, and the responses to series of blue flashes were recorded. For *RPE65* and *GNAT1*-edited tadpoles, responses to a series of flashes of increasing intensity were recorded to measure light sensitivity. For *SAG* and *GRK1*-edited tadpoles, responses to a series of identical flashes presented at regular intervals following a bright light illumination were recorded to measure dark adaptation.

### Statistics

Statistical tests were performed using SPSS vs 25 software (IBM). For dot blot data, statistical tests were performed following log-transformation of the data.

## Results

### Cyclic light is required for RD associated with T4K rhodopsin

We previously generated *X. laevis* carrying a human *RHO* T4K transgene (Tam and Moritz, 2009), a mutation associated with sector RP in humans (van den Born et al., 1994). We determined that RD can be induced in these animals by exposure to 1700 lux cyclic light (12 hours on, 12 hours off), conditions that do not promote RD in wildtype *X. laevis*. Light-induced toxicity was shown to be dependent on photoactivation of the mutant rhodopsin, since a variant that cannot be activated did not cause RD (Tam and Moritz, 2009). Based on this, we hypothesized that generating large amounts of the activated mutant rhodopsin in a short period of time via synchronized photoactivation would be more toxic than asynchronous photoactivation. We therefore raised tadpoles in a more rapid cycling regimen (1 hour on, 1 hour off) as well as conditions of constant light and constant dark (Figure 1A, A’). The offspring of a cross between a *RHO* T4K transgenic male and a wildtype female (yielding 50% transgenic and 50% non-transgenic animals) were raised in complete darkness for 2 weeks, and then held at the specified lighting condition for a further 7 days, after which the animals were sacrificed, and one eye was processed for dot blot assays for rod opsin, while the other was processed for confocal microscopy. Duplicate dot blots were probed with either monoclonal 1D4 to distinguish transgenic from non-transgenic samples, or monoclonal B630N to determine total rod opsin levels for all samples (Tam and Moritz, 2007).

**Figure 1:**
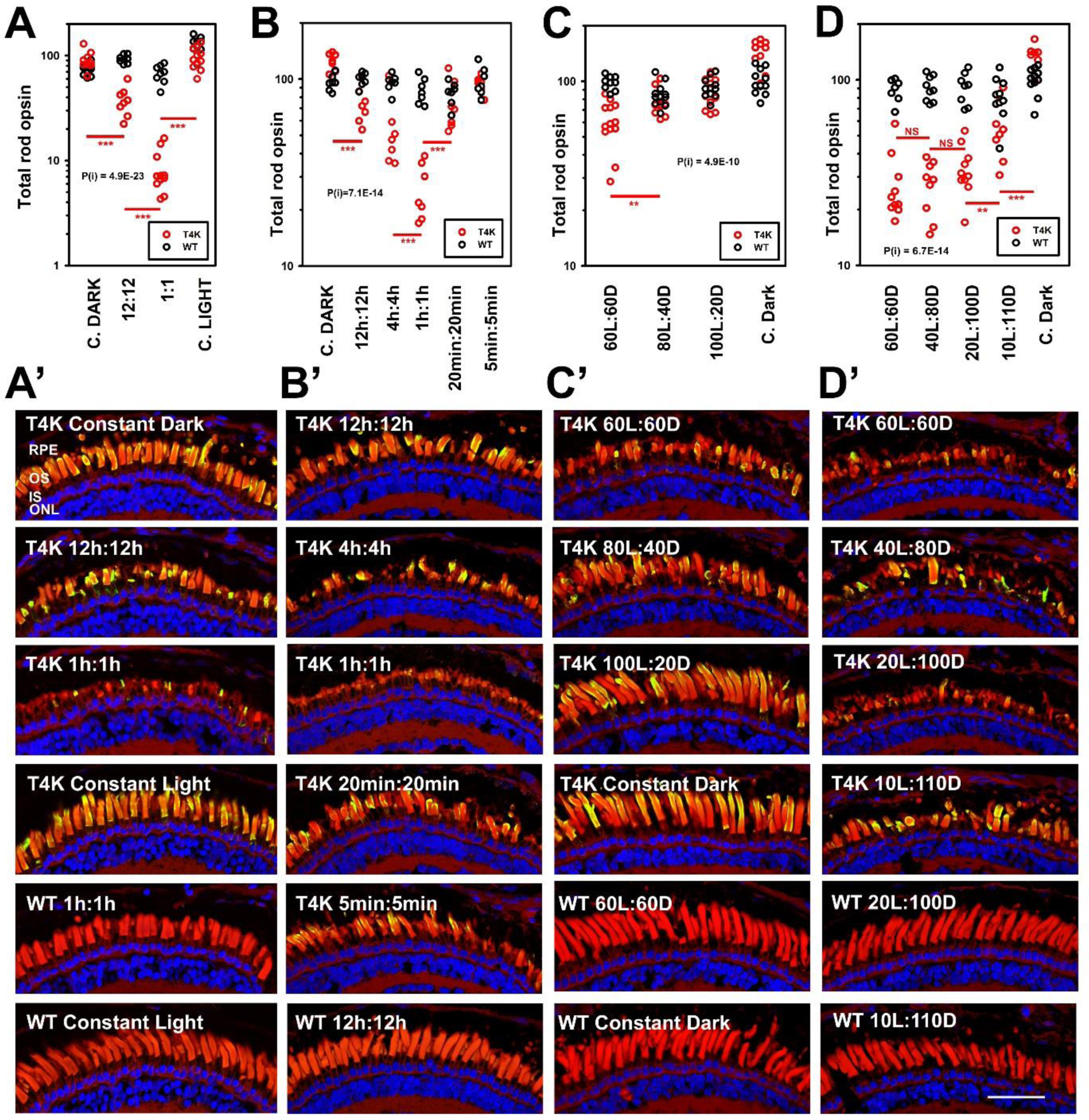
Effects of altered light cycles on RD induced by T4K rhodopsin. **A:** Total rod opsin levels of tadpole eye extracts determined by dot blot analysis using mAbB630N. Tadpoles were reared in darkness to 14 dpf, then transferred to four different lighting regimens of constant darkness, constant light, 12h on 12 hour off, and 1 hour on 1 hour off. Red symbols: T4K rhodopsin transgenics. Black symbols: WT. **B:** A similar experiment with tadpoles reared in constant darkness to 14dpf, then transferred to six different cyclic lighting regimens for 24 hours, and returned to darkness for an additional 24 hours, including constant darkness; 12h:12h off; 4h:4h; 1h:1h; 20min:20min; and 5min:5min, and then returned to darkness for 24 hours. **C:** A similar experiment with tadpoles reared in constant darkness to 12dpf, then transferred to four different lighting regiments of 60 min light, 60 min dark; 80 min light, 40 minutes dark; 100 min light, 20 minutes dark, and constant dark, and then returned to darkness for 24 hours. **D:** A similar experiment with tadpoles reared in constant darkness to 12dpf, then transferred to 5 different cyclic lighting conditions of 60 min light, 60 min dark; 40 min light, 40 min dark; 20 min light, 20 min dark; 10 min light, 10 min dark, and constant dark, and then returned to darkness for 24 hours. Results for statistical test for interaction of genotype and rearing conditions by 2-way ANOVA are shown on each plot. P values for 2-way ANOVA and interaction between condition and genotype were highly significant: A: P = 2.6E-39, P(i) = 4.9E-23; B: P = 1.7E-22, P(i) = 7.1E-14; C: P = 9.2E-16, P(i) = 4.9E-10; D: P = 2.3E-27, P(i) = 6.7E-14. Red bars indicate Tukey post-hoc comparisons between *RHO* T4K groups. Results for selected post-hoc tests are shown as asterisks: * < 0.05, ** < 0.01, *** < 0.001. **A’-E’**: Confocal microscopy of cryosections from representative contralateral eyes of animals described in A-D. Sections are labeled with mAb2B2 (anti-mammalian rhodopsin) (green), WGA (red), and Hoechst33342 (blue).

In agreement with our previous studies(Tam and Moritz, 2009; Tam et al., 2014), relative to their WT siblings, *RHO* T4K transgenic animals exhibited reduced total rod opsin levels (indicative of RD) when reared in standard 12 hour on: 12 hour off cyclic light (Figure 1A), but not when reared in constant darkness. The loss of total rod opsin was more pronounced in transgenic animals exposed to the rapid 1 hour on: 1 hour off cycling regimen, even though the total amount of light exposure was identical. Moreover, there was minimal loss of rod opsin in transgenic animals when they were subjected to constant light, even though they received twice the total light exposure of either cyclic light condition. Confocal images of the cryosectioned contralateral eyes confirmed that reduced total rod opsin was due to loss of rod outer segments (Figure 1A’).

Our results therefore demonstrate that the frequency of the light cycle has a more significant influence on this RD than total light exposure time, and that synchronized activation and regeneration of mutant rhodopsin causes RD, while asynchronous activation does not.

### Optimizing the light cycling regimen for maximal retinal toxicity

In order to obtain additional insight into the underlying mechanisms of RD, we characterized the parameters of the light and dark period required to produce maximal toxicity. We further varied the length of the cycles to include 12 hour, 4 hour, 1 hour, 20 minute, and 5 minute alternating light and dark periods. Cycling was carried out for 24 hours, after which the animals were returned to darkness for 24 hours to allow for clearance of rod outer segment debris before eyes were enucleated. The most toxic condition determined by dot blot assay, and subsequently confirmed by confocal microscopy, was 1 hour on, 1 hour off (i.e. 12 cycles per 24 hours) (Figure 1 B, B’), with minimal RD occurring in the 5 minute on, 5 minute off condition (i.e. 72 cycles per 24 hours), suggesting that this closely resembled the condition of constant light in Figure 1A.

In order to determine whether the reduced RD at the most rapid light cycles was due to insufficiently long periods of darkness, or insufficiently long periods of light, we reared animals under conditions in which the number of light cycles was held constant at 12 cycles per 24 hours, and the period of darkness was decreased while the period of light was proportionately increased. We found that decreasing the period of darkness to 40 minutes or less prevented RD (Figure 1C, C’). Similarly, we decreased the period of light while increasing the period of darkness, and found that reducing the light exposure from 20 to 10 minutes resulted in decreased RD (Figure 1D, D’). Collectively, the results indicate that maximal light-induced RD requires a minimum of 20 minutes light exposure and one hour of regeneration per light cycle.

### The time course of retinal degeneration caused by rapidly cycling light

To characterize the time course of RD in *RHO* T4K transgenic *X. laevis* exposed to rapidly cycling light, we raised F1 transgenic tadpoles and their wild type siblings in complete darkness until they reached 14 dpf, and then transferred them to a 1 hour on: 1 hour off lighting regimen. Animals were sacrificed at specific timepoints just prior to the end of a “lights on” period, and were analyzed by dot blot of total rod opsin, genotyping, and confocal microscopy. Under these conditions, the effects of light on *RHO* T4K transgenics were apparent by 25 hours, and virtually complete by 59 hours. This implies that RD was initiated prior to 25 hours, the timepoint where loss of rod opsin was first apparent in the dot blot assay (Figure 2A and B).

**Figure 2:**
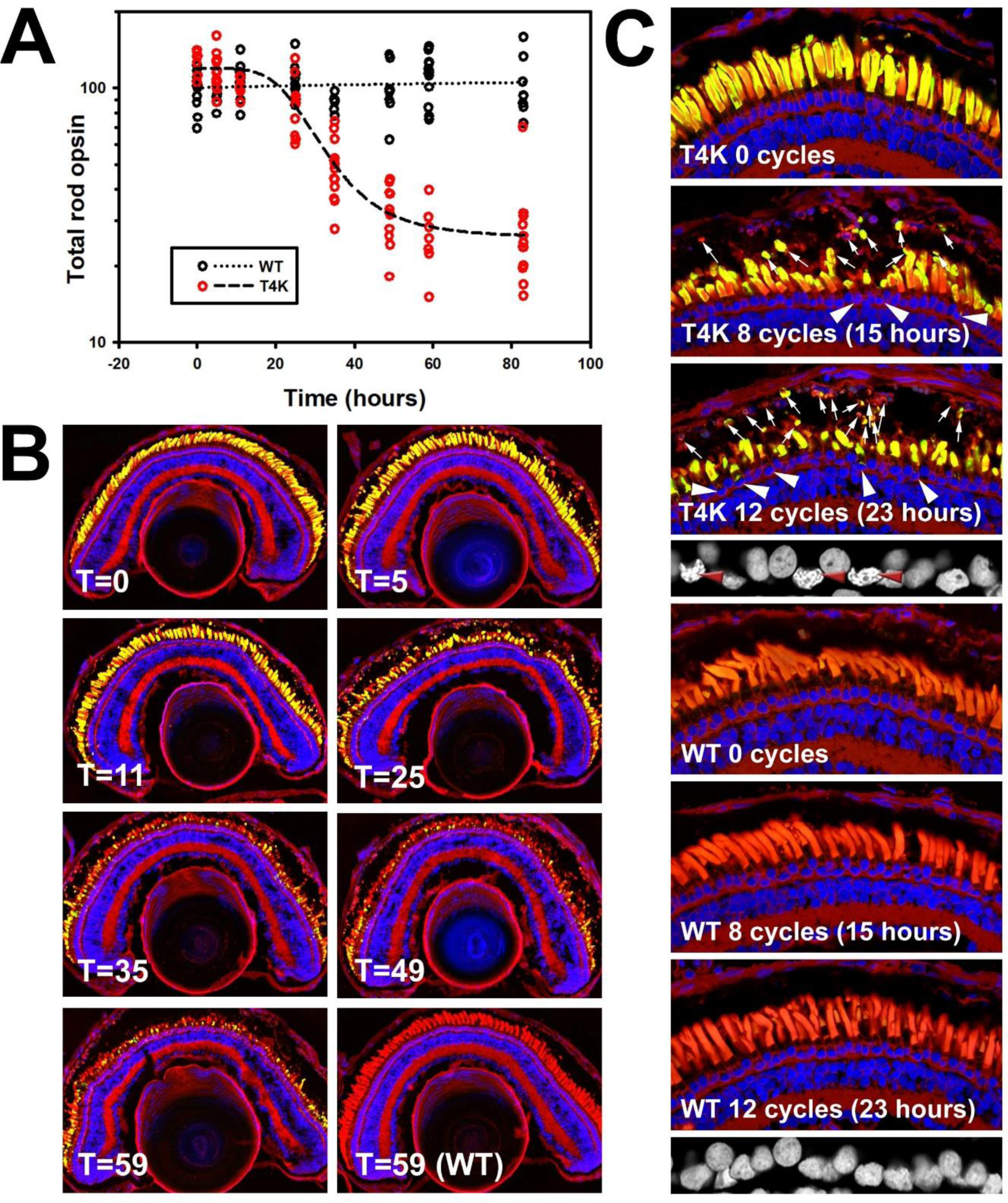
Timecourse of RD in *RHO* T4K animals induced by rapidly cycling light. **A:** Total rod opsin levels determined by dot blot analysis using mAbB630N of eye extracts from tadpoles reared in darkness to 14 dpf, then transferred to a 1hr on:1hr off lighting regimen at T=0, with further timepoints as indicated. Black symbols: WT. Red symbols: *RHO* T4K. Each data point represents one animal. **B:** Confocal microscopy of cryosections from representative contralateral eyes of animals described in **A**. Sections are labeled with mAb2B2 (anti-mammalian rhodopsin) (green), WGA (red), and Hoechst33342 (blue). C: Confocal microscopy of samples from a similar experiment with additional timepoints. Labeling is identical to that in B. Arrows show phagosomes. Arrowheads identify nuclei with abnormal morphology, not apparent at this magnification. Greyscale insets show higher magnification of nuclei, red arrowheads similarly identify nuclei with abnormal morphology.

To further capture these early events, we conducted a similar experiment with more timepoints between T0 (light onset) and T23 hours (12 light cycles). Enucleated eyes were processed for confocal microscopy and the contralateral eye for electron microscopy. Retinal cryosections were labeled with anti-mammalian rhodopsin, counterstained with wheat germ agglutinin and Hoechst (Figure 2C). Changes in the retinas were readily apparent by 8 cycles (i.e. 15 hours following transfer from darkness) including massive shedding of ROS fragments into the RPE. An increase in Hoechst-positive material within the RPE layer indicated phagocytosis of entire cell bodies including their nuclei. By 12 cycles (23 hours) the RPE was densely filled with the remnants of cell bodies and ROS, and partial degradation of this material by the RPE likely produced the initial drop in rod opsin levels that we observed in Figure 2A. In addition, some photoreceptor nuclei in the ONL exhibited altered morphologies consistent with pyknosis. None of these features were apparent in the retinas of WT siblings (Figure 2C).

At the level of electron microscopy (Figure 3), at 8 cycles we similarly observed abundant large ROS-derived phagosomes within the RPE, as well as ROS that appeared to be fragmenting and actively undergoing phagocytosis by the RPE (Figure 3A). At higher magnification, ROS morphology appeared relatively normal, with well-ordered disks apparent in cells with optimal orientation, and even within ROS fragments undergoing phagocytosis (Figure 3C-C’ and D-D’). In contrast, in light-exposed bovine *RHO* P23H animals ROS membranes are markedly disorganized (Figure 3G) as previously reported (Haeri and Knox, 2012; Bogéa et al., 2015).

**Figure 3:**
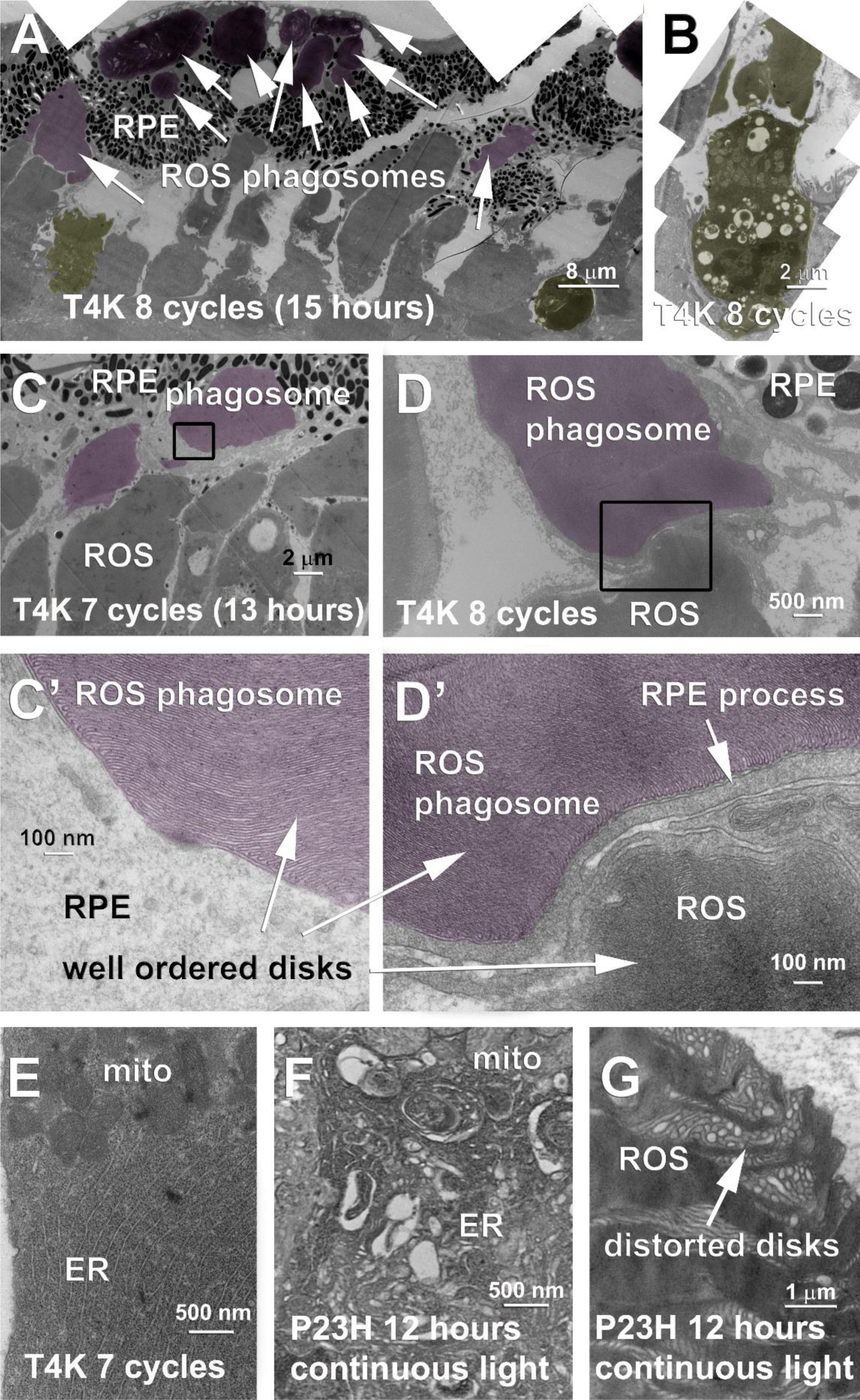
Transmission electron microscopy of *RHO* T4K retinas exposed to rapid cycling. **A:** Overview of a retina exposed to 8 cycles of 1 hr on: 1 hr off lighting. Numerous ROS phagosomes are apparent (arrows, tinted magenta), as well as cells with abnormal nuclear morphology and cytoplasmic vacuolization (tinted yellow). **B:** higher magnification image of a cell with abnormal nuclear morphology and cytoplasmic vacuolization (tinted yellow). **C, D** ROS fragments undergoing phagocytosis from the RPE, taken from 7- and 8-cycle timepoints. Phagosomes are tinted magenta. **C’, D’**: higher magnification images of the areas indicated in C and D. ROS disks are visible, and do not appear disorganized. **E**: Higher magnification image of a T4K transgenic rod inner segment from a 7-cycle timepoint. The membranes of the ER and mitochondria are well organized. **F, G:** Contrasting results from samples expressing bovine P23H rhodopsin and exposed to 12 hours of continuous light. Inner segment membranes are disarrayed and contain autophagic structures. Outer segment disks are distorted and disorganized.

Darkly-staining cells with altered nuclear morphologies and large vacuoles were present within the ONL (Figure 3A and B). In cells without altered nuclear morphology, inner segment appearance was relatively normal, including well-organized ER membranes (Figure 3E). A general absence of intermediate forms suggests that alteration of nuclear morphology and vacuolization is a rapid process. This is in contrast to the altered inner segment morphology we have previously observed in animals carrying a bovine *RHO* P23H transgene reared in the dark and then exposed to light, in which ER membranes are less organized, and autophagic structures are abundant (Bogéa et al., 2015) (Figure 3F). Altogether these results suggest that light induced toxicity originated in the outer segments of T4K rhodopsin-expressing rods and that the downstream phenotype is significantly different from that of P23H rhodopsin-expressing rods.

### Strategies for minimization of RD caused by the interaction of T4K rhodopsin and cyclic light

Frequent cycling of light (1:1) was highly toxic to T4K rhodopsin-expressing retinas, while reducing the frequency of cycles reduced the extent of RD, and constant light had a virtually complete protective effect. We therefore hypothesized that synchronous activation of a threshold number of mutant rhodopsins is required to induce RD, and that other means of reducing synchronous activation would produce protection similar to constant light.

We therefore attempted to attenuate RD in animals expressing T4K rhodopsin by reducing the intensity of cyclic light. Transgenic tadpoles and their wild type siblings were raised in constant dark for 12 days after which they were exposed to 15 cycles (32 hours) of various light intensities on a 1hour on:1 hour off regimen and sacrificed 24 hours after the end of light exposure. Quantitation of total rod opsin and immunofluorescence microscopy demonstrated loss of rod outer segments in transgenic retinas compared to their wild type siblings at light exposures between 1900 lux and 750 lux. The extent of RD was indistinguishable at 1900 and 1475 lux, significantly reduced at 750 lux, and largely eliminated at 370 lux (Figure 4A, D).

**Figure 4:**
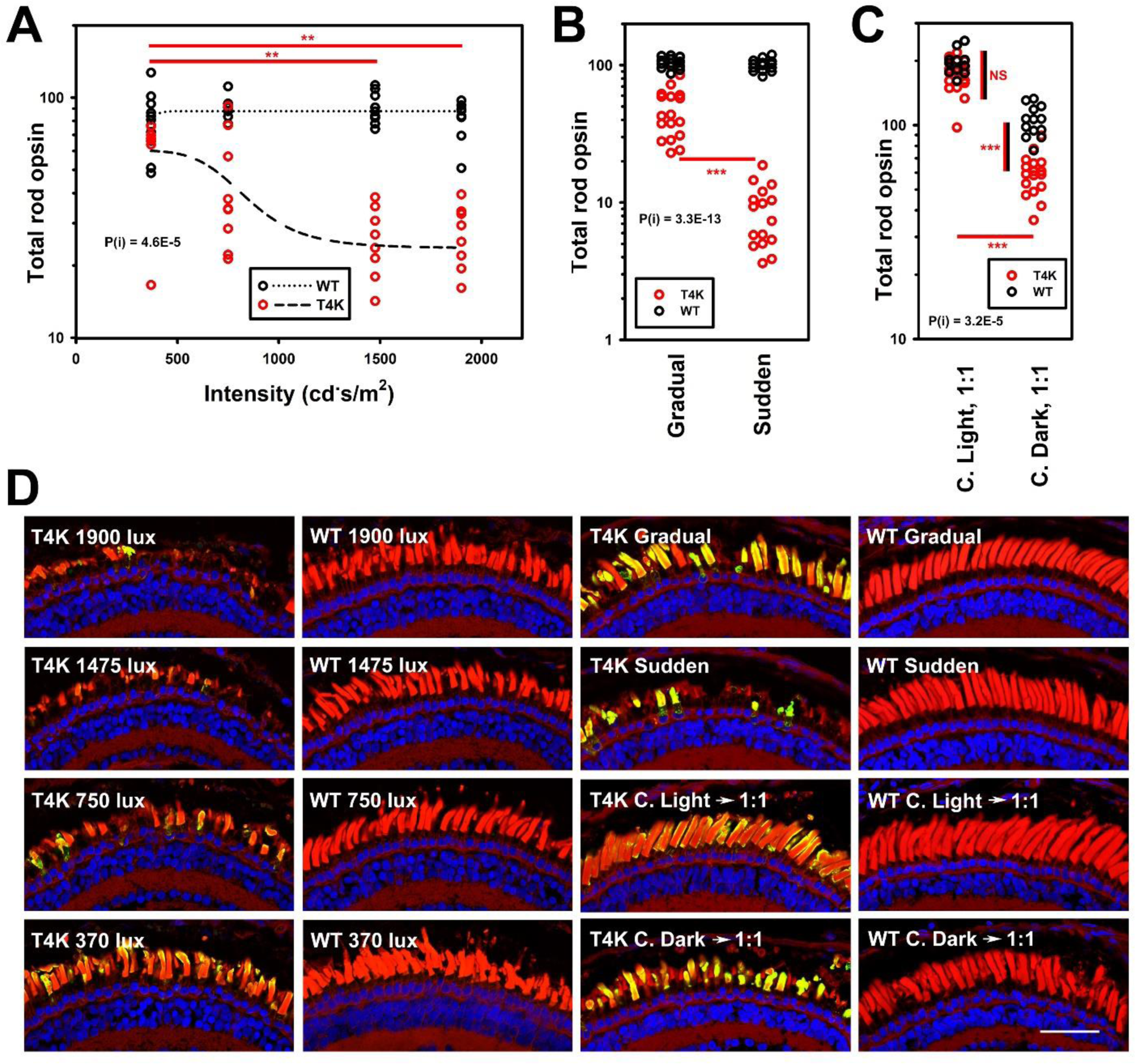
Lighting regimens that alter RD caused by cyclic light in *RHO* T4K retinas. **A:** Total rod opsin levels determined by dot blot analysis using mAbB630N of eye extracts from tadpoles reared in a 12 hr on:12 hr off lighting regimen from 2 dpf to 9 dpf. Light intensity was varied as shown on the plot. Black symbols: WT. Red symbols: *RHO* T4K. Each data point represents one animal. **B:** Similar experiment in which tadpoles were subjected to gradual light onset, or standard 12 hr on: 12 hr off lighting regimen. **C:** Similar experiment in which tadpoles were reared in darkness to 12 dpf, then transferred to either constant light or constant darkness for 24 hours, transferred to rapidly cycling light for 24 hours, and then darkness for a subsequent 48 hours. P values for 2-way ANOVA and interaction between condition and genotype were highly significant: **A:** P = 3.4E-16, P(i) = 4.6E-05; **B:** P = 2.3E-29, P(i) =3.3E-13; **C:** P = 3.5E-34, P(i) = 2.2E-06. **D:** Confocal microscopy of cryosections from representative contralateral eyes of animals described in **A-C.** Green: mAb2B2 anti-mammalian rhodopsin. Red: WGA. Blue: Hoechst33342. **A-C:** Results for statistical test for interaction of genotype and rearing conditions by 2-way ANOVA are shown on each plot. Results for selected Tukey post-hoc tests are shown as asterisks: * < 0.05, ** < 0.01, *** < 0.001. Red lines indicate post hoc tests comparing groups of *RHO* T4K animals. Red and black lines indicate post hoc tests comparing groups of *RHO* T4K and WT animals.

Our standard cyclic lighting regimens involve abrupt transition between dark and light, resulting in a synchronized burst of rhodopsin activation, after which activation would occur asynchronously at a rate limited by chromophore regeneration. Therefore, we hypothesized that eliminating sudden light onset might also reduce RD. Transgenic tadpoles and their wild type siblings were reared for seven days in 12 hours on: 12 hours off with either abrupt or gradual dark-to-light transitions. For gradual transitions, at the beginning of the light phase, lights were turned on at low intensity, and intensity was successively doubled in 10 steps over a 2-hour period until reaching maximum intensity (i.e. 2-hour ramp, 12 hours on, 10 hours off). We found that sudden onset was considerably more toxic to photoreceptors expressing T4K rhodopsin than gradual onset (Figure 4B, D), even though gradual onset involved two additional hours of light exposure.

We have previously shown that *in vitro*, T4K rhodopsin is defective in its ability to regenerate (Tam et al., 2014). We therefore hypothesized that *in vivo*, T4K rhodopsin may be limited to a finite number of activation/regeneration cycles. Therefore, we investigated whether pre-treatment with constant light would protect against light-induced RD. We reared animals in constant darkness or constant light for 12 days post-fertilization, and then subjected them to a rapid light cycling protocol (1 hour on: 1 hour off) for 24hrs, followed by a 24hr period in the dark to allow clearance of phagocytosed. Animals reared in constant darkness followed by rapidly cycling light had significant loss of rod opsin and RD relative to their wildtype siblings.

In contrast, animals reared in constant light followed by rapidly cycling were not significantly different from their WT siblings (Figure 4C, D). Constant light-pretreated animals were larger than the constant dark-pretreated animals and thus had higher rod opsin levels. However, as the critical comparison is between transgenic and wild type within each group, our conclusion is unaffected. Altogether, our results indicate that decreased light intensity, gradual light onset, and pre-treatment with constant light were all protective against RD caused by T4K rhodopsin.

### Genetic interaction of *RPE65* and *RHO* T4K]

Previously we determined that dietary deprivation of vitamin A can prevent light-induced RD caused by T4K rhodopsin, as does elimination of the chromophore binding site (Tam et al., 2014). We hypothesized that this was due to the essential role of the vitamin A-derived 11-cis retinal as the chromophore required for rhodopsin activation, and that therefore we should be able to produce a similar effect by disruption of the *RPE65* gene, which encodes the isomerohydrolase that is critical for regeneration of 11-cis retinal from all-trans retinal in the visual cycle (Redmond et al., 1998; Jin et al., 2005). To test this hypothesis, we used a paradigm based on gene editing with CRISPR-Cas9 to abolish RPE65 expression/activity. We designed an sgRNA directed at a conserved sequence found in both *X. laevis RPE65* genes (*RPE65.S* and *RPE65.L*) and co-injected the sgRNA and Cas9 mRNA into wildtype single-cell embryos. At 14 days post-fertilization we examined the resulting animals by western blot of solubilized eyes. We found that co-injection of sgRNA and Cas9 mRNA largely eliminated expression of the RPE65 protein (Figure 5A and B). In addition, Sanger sequencing analysis of PCR products derived from the *RPE65* genes showed evidence of editing at the predicted cleavage sites (Figure 5A). We examined the scotopic electroretinogram (ERG) of these tadpoles. Consistent with previous studies in mice (Redmond et al., 1998) we found that the ERG was severely compromised (Figure 5C and D). Collectively, these results are consistent with successful disruption of the *X. laevis RPE65* genes.

**Figure 5:**
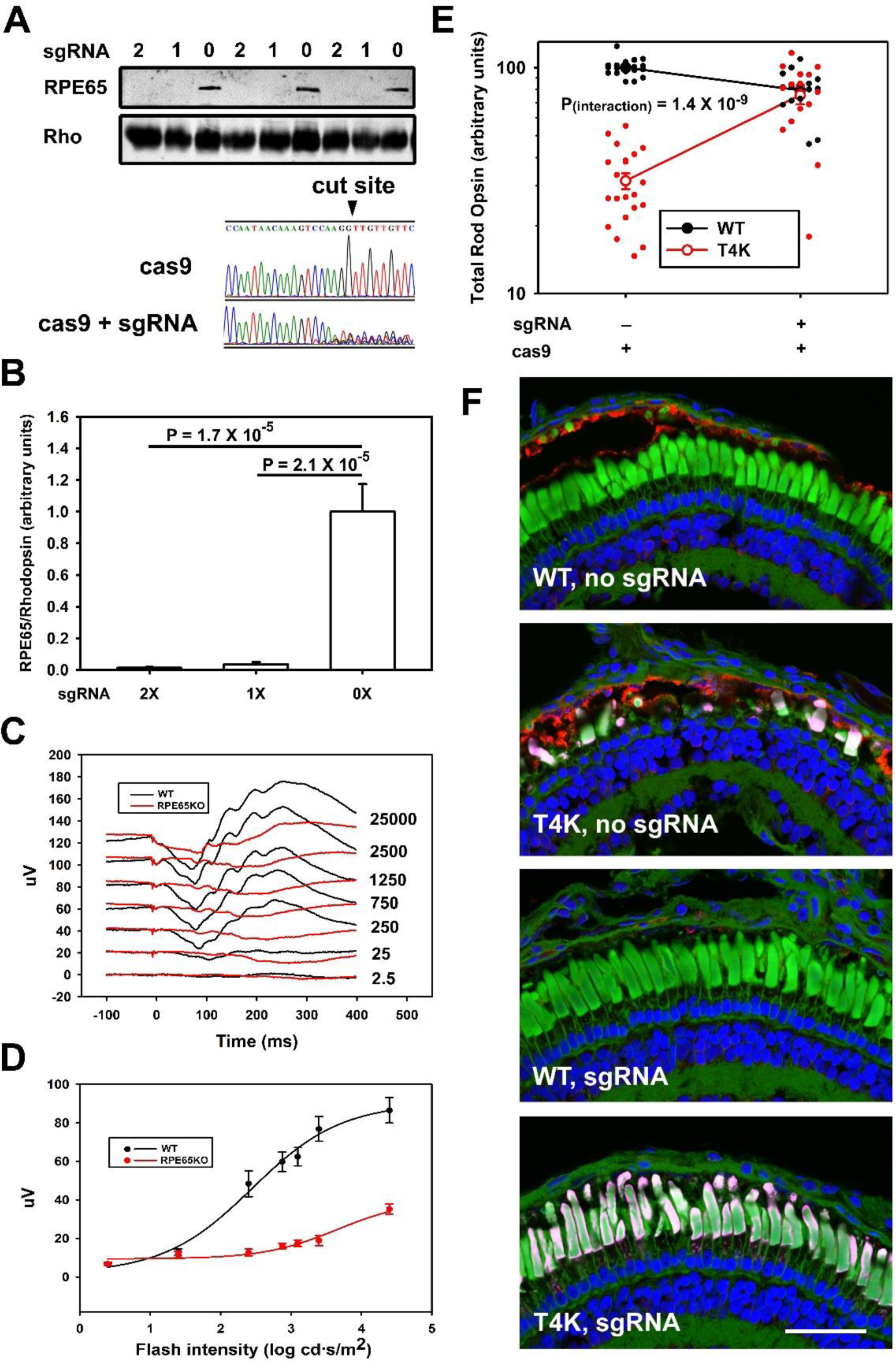
Genetic ablation of RPE65, and genetic interaction of *RPE65* and *RHO* T4K. **A**, **top:** western blot of solubilized eyes from 14dpf animals injected with 0, 1, or 2 units of an sgRNA targeting the *RPE65* genes and Cas9 mRNA, detected with anti-RPE65 or mabB630N anti-rod opsin. **A, bottom:** Direct Sanger sequencing of a PCR product derived from the *RPE65.L* gene from edited and non-edited embryos, encompassing the predicted editing site (indicated). **B:** quantification of the anti-RPE65 western blot shown in **A**, normalized to the anti-rhodopsin signal. **C:** averaged electroretinography of *RPE65*-edited animals (red, n = 7) and unedited WT controls (black, n = 5). **D:** B-wave analysis of data shown in **C**. **E:** dot blot analysis of solubilized eyes from *RHO* T4K animals and their WT siblings, injected at the one-cell stage with RPE65-targeting sgRNA and Cas9, or Cas9 only. Each data point represents an independently derived animal. Mean values +/- S.E.M. are also shown. Results were analyzed by 2-way ANOVA: P = 1.7 × 10^−17^, P_(interaction)_ = 1.4 × 10^−9^. **F:** Confocal microscopy of cryosections from representative contralateral eyes of animals described in **E.** Ablation of RPE65 prevents the retinal degeneration caused by *RHO* T4K. Red: anti-RPE65. Magenta: anti-mammalian rhodopsin mAb2B2. Green: WGA. Blue: Hoechst33342.

Subsequently, we injected the *RPE65*-targeting sgRNA into single-cell embryos derived from a cross between WT and *RHO* T4K transgenic animals, from which half the resulting animals were WT and half carried the *RHO* T4K transgene. Embryos were either injected with sgRNA + Cas9 mRNA or with Cas9 mRNA only, to create four conditions in total. We reared the animals in standard 12 hour on: 12 hour off cyclic light for 14 days, and then performed our standard rod opsin dot-blot and genotyping assays, as well as confocal microscopy using anti-mammalian rod opsin and anti-RPE65 antibodies for immunolabeling. In WT animals, sgRNA resulted in ablation of RPE65 expression as seen by reduction of labeling (green) compared to animals that did not receive sgRNA (Figure 5F). Lack of RPE65 caused a small reduction in rod opsin levels with little or no RD (Figure 5E and F). As expected, *RHO* T4K animals exhibited extensive RD accompanied by loss of total opsin. Strikingly however, ablation of RPE65 expression resulted in virtually complete prevention of RD. In T4K animals that received the sgRNA, rod opsin was at WT levels and histology was normal. The interaction between genotype and treatment was highly significant by 2-way ANOVA (P interaction = 1.4 × 10^−9^). Overall, the results indicate that disruption of the *RPE65* genes prevents RD caused by T4K rhodopsin.

### Genetic interaction of *RPE65* and *RHO* P23H

We have previously found that dietary deprivation of vitamin A promotes RD in *X. laevis* expressing P23H rhodopsin (Tam et al., 2010), likely due to a pharmacological chaperone effect of the chromophore that stabilizes the P23H protein during biosynthesis. Therefore, we hypothesized that disruption of the *RPE65* genes would promote RD in *RHO* P23H transgenic animals, unlike the protective effect seen in *RHO* T4K transgenics. We used CRISPR/Cas9 to edit the *RPE65* genes in animals expressing bovine P23H rhodopsin and determined the consequences of the loss of RPE65 function in animals reared in either 14 days of darkness or cyclic light.

As expected, P23H rhodopsin-expressing animals reared in cyclic light exhibited extensive RD compared to their wild type siblings (Figure 6B). This was also true for edited animals that were injected with sgRNA+cas9 mRNA. The reduction in rod opsin levels in *RHO* P23H transgenics (Figure 6A) was similar for both injected and uninjected groups indicating that loss of RPE65 activity had minimal effect on RD. There was however a slight reduction in the amount of rod opsin in sgRNA+cas9 mRNA injected wildtype animals (Figure 6A), likely due to reduced efficiency of biosynthesis in the absence of chromophore, which led to a small but significant interaction between genotype and treatment.

**Figure 6:**
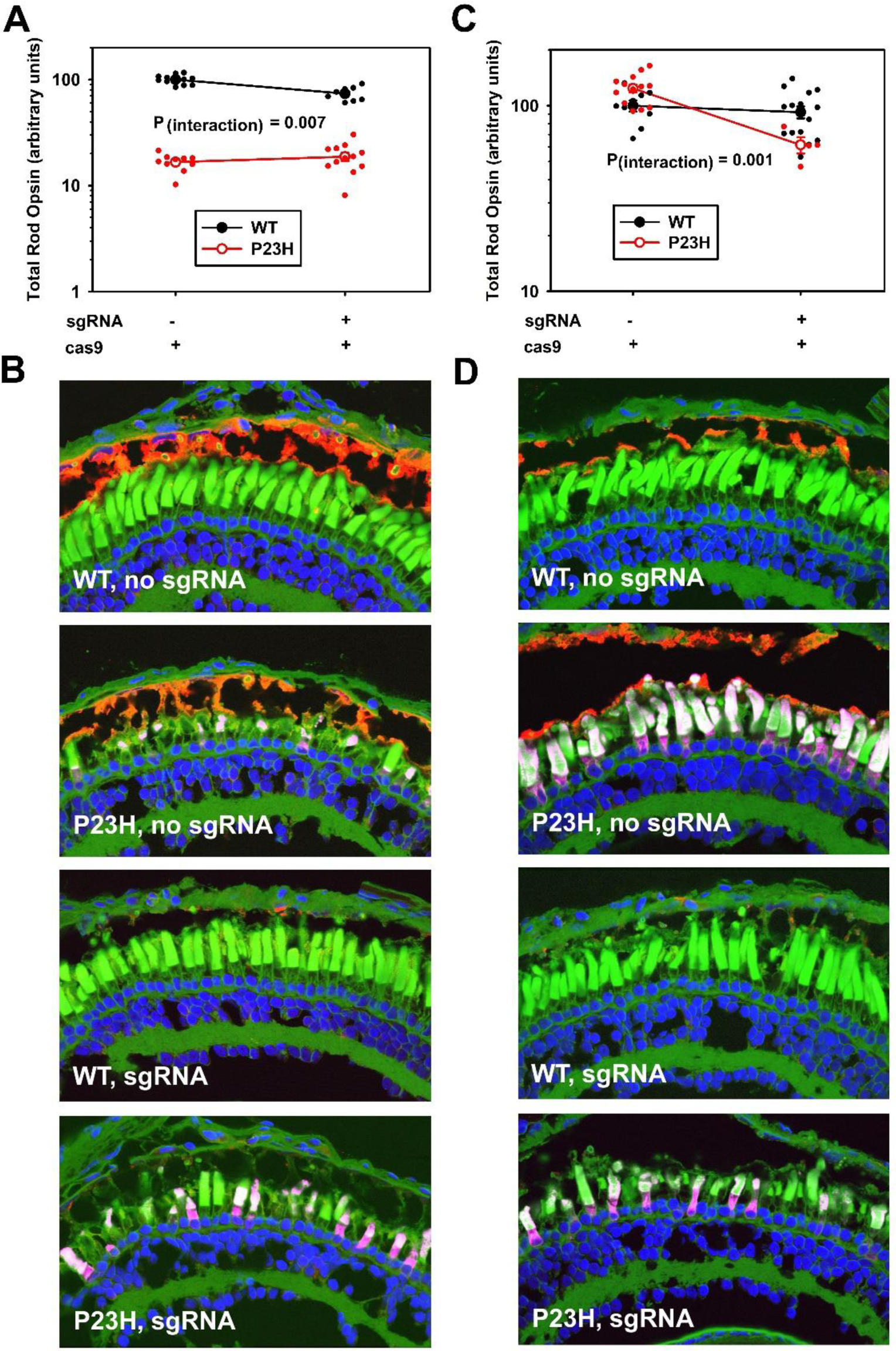
Genetic ablation of RPE65, and genetic interaction of *RPE65* and *RHO* P23H. **A:** Total rod opsin levels determined by dot blot analysis using mAbB630N for solubilized eyes from bP23H rhodopsin transgenic animals and their WT siblings, reared to 14dpf in standard cyclic light, and injected at the one-cell stage with RPE65-targeting sgRNA and Cas9 mRNA, or Cas9 mRNA only. Each data point represents an independently derived animal. Mean values +/- S.E.M. are also shown. Results were analyzed by 2-way ANOVA: P = 5.0E-23, P_(interaction)_ = 6.9E-03. **B:** Confocal microscopy of cryosections from representative contralateral eyes of animals described in **A. C:** experiment identical to that shown in A, but animals were reared in darkness to 14dpf. Results were analyzed by 2-way ANOVA: P = 6.5E-05, P_(interaction)_ = 1.1E-03. **D:** Confocal microscopy of cryosections from representative contralateral eyes of animals described in **C.** Ablation of RPE65 promotes RD in dark reared *RHO* P23H animals. Red: anti-RPE65. Magenta: anti-mammalian rhodopsin mAb2B2. Green: WGA. Blue: Hoechst33342.

Dark rearing of unedited *RHO* P23H animals largely protected against RD as previously seen(Tam and Moritz, 2007). However *RHO* P23H animals with ablated RPE65 expression exhibited reduced rod opsin accompanied by RD (Figure 6C and D), resulting in a significant interaction effect by 2-way ANOVA (P = 0.001). Thus, elimination of RPE65 did not ameliorate RD in cyclic light but did result in loss of protection conferred by dark rearing. Altogether, these results demonstrate that 11-cis retinal has opposing effects on RD caused by P23H rhodopsin versus T4K rhodopsin. The relative lack of rescue from P23H rhodopsin toxicity was in stark contrast to the complete rescue observed for T4K rhodopsin under the same conditions.

### Genetic interaction of *GNAT1* and *RHO* T4K

Since RD caused by T4K rhodopsin requires photoactivation of the mutant rhodopsin, we hypothesized that downstream phototransduction cascade partners might also participate in the RD mechanism. We therefore applied our CRISPR methodology for probing genetic interactions with mutant *RHO* transgenes to *GNAT1*, which encodes rod alpha-transducin, the G-protein activated by rhodopsin. We designed two sgRNAs (sg1 and sg2) that each targeted both *X. laevis GNAT1* genes (*GNAT1.S* and *GNAT1.L*). We co-injected these sgRNAs with Cas9 mRNA and analyzed GNAT1 expression levels by western blot. A random sgRNA (sgN) was included as a control. Relative to injection with sgN, injection with the *GNAT1*-directed sgRNAs promoted loss of the GNAT1 protein (Figure 7A and B), and caused *GNAT1* gene editing detectable by Sanger sequencing (Figure 7A). Sg2 was more effective at reducing gene expression and was therefore used in subsequent experiments. To demonstrate the effect of *GNAT1* editing on function, scotopic ERGs were performed. Disruption of *GNAT1* severely compromised visual function of edited animals (Figure 7C and D). Loss of scotopic ERG due to targeted deletion of *GNAT1* has also been observed in mice (Calvert et al., 2000).

**Figure 7:**
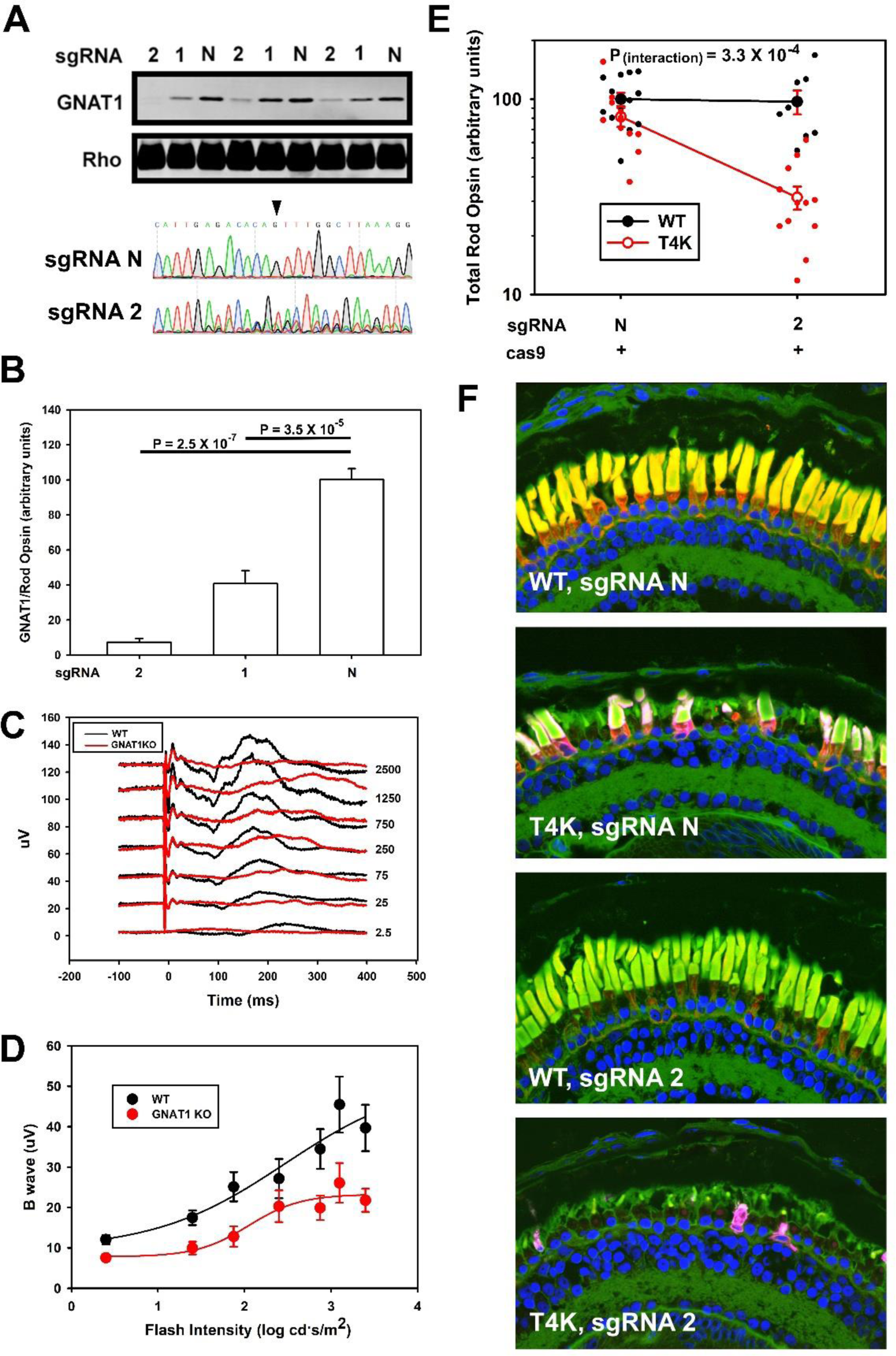
Genetic ablation of GNAT1, and genetic interaction of *GNAT1* and *RHO* T4K. **A**, **top:** western blot of solubilized eyes from 14dpf animals injected with a Cas9 mRNA and either control sgRNA “N” or equal quantities of two different sgRNAs targeting the *GNAT1* genes (1 and 2), detected using anti-GNAT1 or mAbB630N anti-rod opsin. **A, bottom:** Direct Sanger sequencing of a PCR product derived from the *GNAT1.L* gene from embryos injected with sgRNA “N” or *GNAT1*-targeting sgRNA2, encompassing the predicted editing site (indicated). **B:** quantification of the anti-GNAT1 western blot shown in **A**, normalized to the anti-rhodopsin signal. P-values are Tukey multimple comparisons tests following ANOVA (P = 3.2 E-07). **C:** averaged electroretinography of *GNAT1*-edited animals (red, n = 9) and controls injected with sgRNA “N” (black, n = 8). **D:** B-wave analysis of data shown in **C**. **E:** Anti-rod opsin dot blot analysis of solubilized eyes from *RHO* T4K animals and their WT siblings, injected at the one-cell stage with *GNAT1*-targeting sg2 and Cas9 mRNA, or Cas9 mRNA and sgN. Each data point represents an independently derived animal. Mean values +/- S.E.M. are also shown. Results were analyzed by 2-way ANOVA: P = 2.9 × 10^−9^, P_(interaction)_ = 3.3 × 10^−4^. **F:** Confocal microscopy of cryosections from representative contralateral eyes of animals described in **E.** Ablation of GNAT1 exacerbates the retinal degeneration caused by *RHO* T4K. Green: anti-GNAT1. Magenta: anti-mammalian rhodopsin mAb2B2. Red: WGA. Blue: Hoechst33342.

After identifying an sgRNA that efficiently disrupted *X. laevis GNAT1*, we looked for a genetic interaction between *GNAT1* and *RHO* T4K in the RD phenotype, similar to the experiment conducted on *RPE65*. We found that disruption of *GNAT1* dramatically reduced rod opsin levels and exacerbated RD in *RHO* T4K transgenics, while causing little or no difference in rod opsin levels in WT animals (Figure 7E and F) (P interaction = 3.3 × 10^−4^). The expression level of GNAT1 was markedly reduced as determined by immunolabeling (Figure 7F). Overall the results indicate that disruption of *GNAT1* exacerbates RD caused by T4K rhodopsin.

### Genetic interaction of *SAG* and *RHO* T4K

We next investigated genetic interactions of *RHO* T4K and *SAG*, the gene encoding rod arrestin, which directly binds phosphorylated rhodopsin in an early event in the shut-off of the phototransduction cascade. We designed two sgRNAs, (sg3 and sg4) to edit the *X. laevis SAG.S* and *SAG.L* genes respectively. We confirmed that simultaneous co-injection of both sgRNAs with cas9 mRNA caused editing of the *SAG* genes, reducing SAG expression by >80% on average, relative to injection with Cas9 mRNA alone (Figure 8A and B). Disruption of *SAG* causes congenital stationary night blindness due to delayed adaptation of photoreceptors to darkness (Fuchs et al., 1995) Therefore, we assessed the effects of *SAG* disruption on dark adaptation using a modified ERG protocol in which we monitored the recovery of the ERG over time following exposure to a bright light. In wild type animals, b-wave amplitudes progressively increased between 30 and 350 seconds after an intial bright light flash. In contrast, SAG-edited animals had a small b-wave which did not recover over the course of the experiment. These results are consistent with disruption of dark adaptation in these animals (Figure 8C and D).

**Figure 8:**
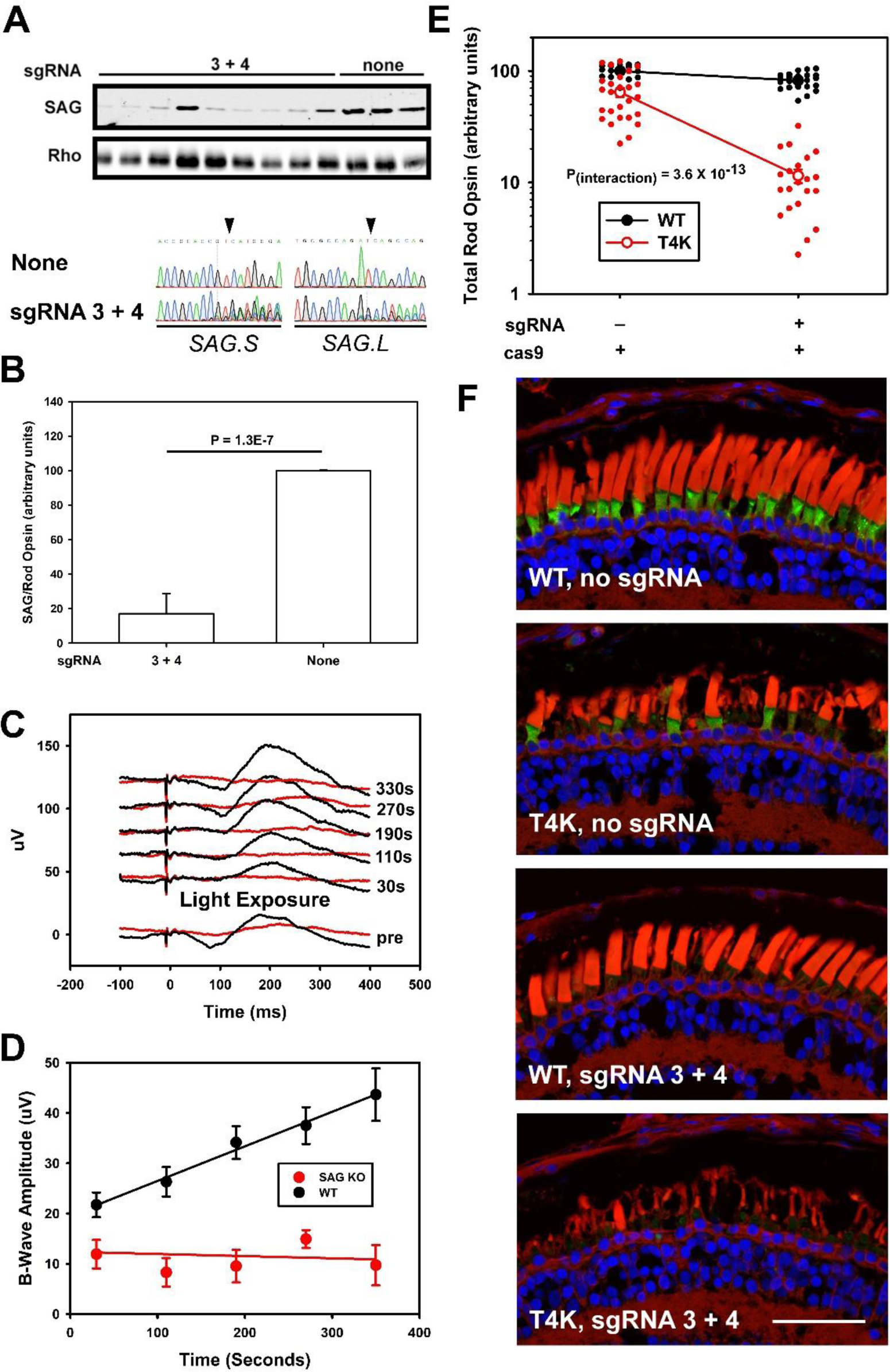
Genetic ablation of SAG, and genetic interaction of *SAG* and *RHO* T4K. **A**, **top:** western blot of solubilized eyes from 14dpf animals injected with Cas9 mRNA or Cas9 mRNA and a combination of two different sgRNAs targeting *SAG.S* and *SAG.L*, detected using mAb xArr1-6 anti-SAG or mAbB630N anti-rod opsin. **A, bottom:** Direct Sanger sequencing of PCR products derived from the *SAG.L* and *SAG.S* genes from edited and non-edited embryos, encompassing the predicted editing site (indicated). **B:** quantification of the anti-SAG western blot shown in **A**, normalized to the anti-rhodopsin signal. P value is derived from T-test. **C:** averaged electroretinography of *SAG*-edited animals (red, n = 8) and WT controls (black, n = 10). An initial response to a blue flash was recorded, followed by a bright light exposure, and then subsequent blue flashes at the specified intervals, to monitor dark adaptation. **D:** B-wave analysis of data shown in **C**. **E:** Anti-rod opsin dot blot analysis of solubilized eyes from *RHO* T4K animals and their WT siblings, injected at the one-cell stage with *SAG-*targeting sgRNAs and Cas9 mRNA, or Cas9 mRNA only. Each data point represents an independently derived animal. Mean values +/- S.E.M. are also shown. Results were analyzed by 2-way ANOVA: P = 3.8 × 10^−31^, P_(interaction)_ = 3.6 × 10^−13^. **F:** Confocal microscopy of cryosections from representative contralateral eyes of animals described in **E.** Ablation of SAG exacerbates the retinal degeneration caused by *RHO* T4K. Green: mAb xArr1-6 anti-SAG. Red: WGA. Blue: Hoechst33342.

We looked for a genetic interaction between *SAG* and *RHO* T4K in the RD phenotype, similar to the experiments conducted on *RPE65* and *GNAT1*. Surprisingly, disruption of *SAG* also dramatically reduced rod opsin levels and exacerbated RD in T4K animals, while causing little or no difference in rod opsin levels in wildtype animals (Figure 8E and F). Confocal micrographs were consistent with loss of SAG expression in the majority of rod photoreceptors (Figure 8F). Thus, RD was exacerbated when either SAG or GNAT1 expression was reduced despite the fact that these proteins have opposite roles in phototransduction.

### Genetic interaction of *GRK1* and *RHO* T4K

We next investigated interactions of *RHO* T4K and *GRK1*, the gene encoding rhodopsin kinase, which phosphorylates rhodopsin to promote binding of SAG in the shut-off of the phototransduction cascade. We created two sgRNAs, each designed to edit both the *X. laevis GRK1.S* and *GRK1.L* genes. By western blot, we determined that both sgRNAs were effective at reducing GRK1 expression, and we observed editing of the *GRK1* genes by Sanger sequencing (Figure 9A and B). Subsequent experiments used sg2.

**Figure 9:**
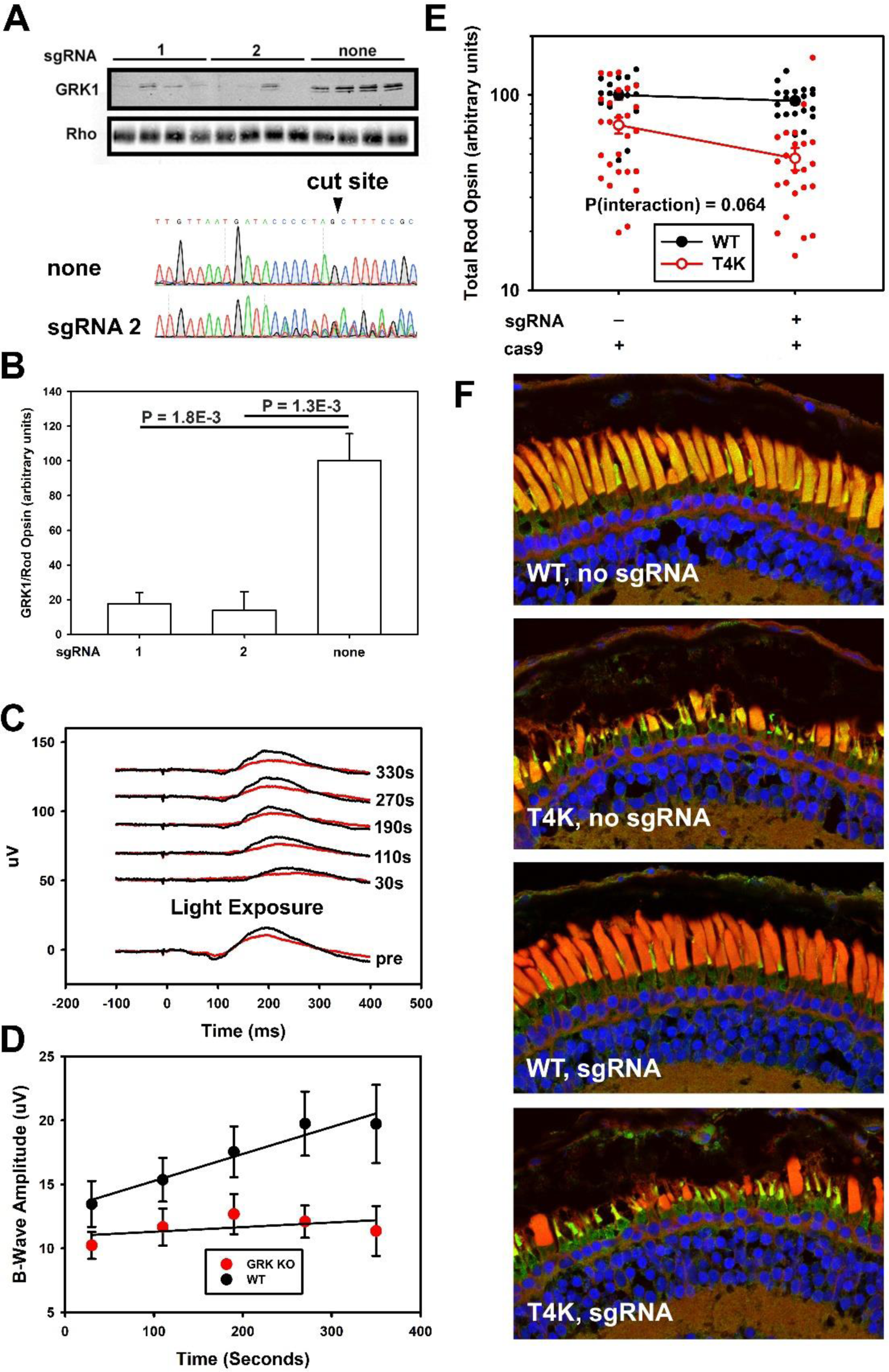
Genetic ablation of GRK1, and genetic interaction of *GRK1* and *RHO* T4K. **A**, **top:** western blot of solubilized eyes from 14dpf animals injected with Cas9 mRNA or Cas9 mRNA and one of two different sgRNAs targeting the *GRK1* genes (1 and 2), detected using mAbG8 anti-GRK1 or mAbB630N anti-rod opsin. **A, bottom:** Direct Sanger sequencing of PCR products derived from the *GRK1.L* gene of non-edited embryos, and embryos edited using *GRK1*-targeting sgRNA2, encompassing the predicted editing site (indicated). **B:** quantification of the anti-SAG western blot shown in **A**, normalized to the anti-rhodopsin signal. P-values are Tukey multimple comparisons tests following ANOVA (P = 7.4 E-04). **C:** averaged electroretinography of *GRK1*-edited animals (red, n = 9) and WT controls (black, n = 8). An initial response to a blue flash was recorded, followed by a bright light exposure, and then subsequent blue flashes at the specified intervals, to monitor dark adaptation. **D:** B-wave analysis of data shown in **C**. **E:** Anti-rod opsin dot blot analysis of solubilized eyes from *RHO* T4K animals and their WT siblings, injected at the one-cell stage with *GRK1-*targeting sgRNA2 and Cas9 mRNA, or Cas9 mRNA only. Each data point represents an independently derived animal. Mean values +/- S.E.M. are also shown. Results were analyzed by 2-way ANOVA: P = 5.6 × 10^−9^, P_(interaction)_ = 0.064. **F:** Confocal microscopy of cryosections from representative contralateral eyes of animals described in **E.** Note that mAbG8 anti-GRK1 cross-reacts with GRK7 expressed in cones. Ablation of GRK1 exacerbates retinal degeneration caused by *RHO* T4K, but the test for interaction of genotype and GRK1 ablation is not statistically significant. Green: anti-GRK1. Red: WGA. Blue: Hoechst33342.

Antibody labeling of retinal sections showed that edited rod OS had reduced or undetectable anti-GRK1 labeling compared to wild type (Figure 9F). However, cone OS in both edited and WT retinal sections showed high levels of labeling, likely due to antibody cross-reactivity with GRK7, the cone-specific kinase. Similar to *SAG*, mutations in *GRK1* cause congenital stationary night blindness (Yamamoto et al., 1997; Chen et al., 1999a) and so we assessed the effect of editing with sg2 on visual function using the same ERG assay used for the *SAG* experiment described above. The results were similar, in that disruption of the *GRK1* genes was associated with delayed dark adaptation (Figure 9C and D).

We looked for a genetic interaction between *GRK1* and *RHO* T4K in the RD phenotype. We found that disruption of *GRK1* reduced rod opsin levels and exacerbated RD in *RHO* T4K transgenics, while causing little or no difference in rod opsin levels in WT animals (Figure 9E and F). However, the 2-way ANOVA test for interaction between treatment and genotype was not significant (P = 0.064), although a 1-way ANOVA followed by Tukey test indicated a significant difference between treated and untreated T4K animals (P = 0.008), and no difference between treated and untreated WT animals (P = 0.982). The reduced statistical significance was due to both a smaller effect and greater variability.

## Discussion

The *RHO* mutation, T4K, is associated with a “sector” or B1” RP phenotype in humans (Bunge et al., 1993; van den Born et al., 1994; Cideciyan et al., 1998). Previous studies indicate that this mutation causes a defect in N-linked glycosylation of rod opsin, and a light-exacerbated RD phenotype that requires photoactivation of the mutant protein (Kaushal et al., 1994; Tam and Moritz, 2009; Tam et al., 2014). Here we show that induction of RD specifically required ***cyclic*** light, as RD was prevented not only by dark-rearing but also by constant light. Moreover, our results suggest that RD is exacerbated by conditions that promote synchronous photoactivation of large numbers of T4K rhodopsin molecules in the ROS (more frequent light cycling, higher light intensities and sudden light onset). RD was reduced under conditions promoting asynchronous photoactivation (reduced light intensity, constant light exposure, pretreatment with light and gradual light onset). Light cycles were most toxic when they consisted of at least one hour of darkness and 20 minutes of light exposure. Genetic ablation of phototransduction binding partners GRK1, SAG and GNAT1, all exacerbated light-induced retinal degeneration associated with T4K rhodopsin, while genetic ablation of RPE65 prevented it. Our results provide further insight into the pathogenic mechanisms involved in this model of RP as well as potential intervention points for modulating the serverity of the disease.

Light exacerbated RD caused by T4K rhodopsin in many respects resembles RDs caused by short intense or prolonged light exposures in wild type mice, rats and zebrafish that are exacerbated by more rapid cycling (NOELL et al., 1966), prevented by loss of RPE65 activity (Grimm et al., 2000; Wenzel et al., 2001), promoted by prolonged dark adaptation (Noell, 1979; Vihtelic and Hyde, 2000), and ameliorated by prior light exposure (Noell, 1979) suggesting possible mechanistic overlaps amongst these models. Moreover, similar to RD induced by *RHO* T4K, RD caused by prolonged light exposure in mice is promoted by disruption of *SAG* and *GRK1* (Chen et al., 1999b, 1999a). Thus, our model of sector RP may represent exaggeration of some aspects of light damage seen in other systems; however, the unique feature of protection by constant light indicates that there are also mechanistic distinctions.

Notably, disruption of *RPE65* expression promotes RD caused by the commonly studied sector RP-associated mutant, *RHO* P23H (Figure 6) but has the opposite effect on *RHO* T4K (Figure 5). Therefore, despite outwardly similar phenotypes, the underlying mechanisms differ and have distinct responses to interventions. While the light-exacerbated RD associated with *RHO* P23H is likely due to loss of the pharmacological chaperone effect of 11-cis retinal (Noorwez et al., 2003; Kosmaoglou et al., 2008; Tam et al., 2010; Chen et al., 2014), the light exacerbated RD associated with *RHO* T4K requires photoactivation of the mutant protein (Tam et al., 2014). We propose distinguishing these mechanisms underlying sector RP as “chromophore-exacerbated RD” (T4K-like) and “chromophore-mitigated RD” (P23H-like). Additionally, both mechanisms could conceivably be present simultaneously in a given mutant (e.g. partial biosynthetic defect and toxic photoproduct).

As chomophore-exacerbated RD requires photoactivated mutant rhodopsin, a reasonable hypothesis is that the pathogenic mechanism may involve prolonged or over-activation of the phototransduction cascade (Fain and Lisman, 1993) as has been proposed in light-induced RD in *SAG* knockout mice (Chen et al., 1999b; Fain and Lisman, 1999) (Hao et al., 2002). Moreover, it has been shown that constitutively active rhodopsins are constitutively bound to SAG, and these complexes can initiate photoreceptor cell death (Alloway et al., 2000; Chuang et al., 2004; Chen et al., 2006; Park, 2014). Because T4K rhodopsin requires photoactivation to induce RD, we hypothesized that similar mechanisms may be involved. However, we found that both inactivation of the phototransduction cascade by disruption of *GNAT1* and prevention of rhodopsin-SAG complex formation by disruption of *SAG* exacerbated the *RHO* T4K RD phenotype, suggesting neither mechanism applies to this model.

As an alternative hypothesis, we propose that the toxic effect is caused by an abnormal photoproduct of T4K rhodopsin. In this scenario, both GNAT1 and SAG ameliorate RD by either preventing adoption of this toxic conformation, preventing its interaction with a target, or facilitating its removal, and neither GNAT1 nor SAG mediates the phototoxic effect. We observed a similar but less statistically significant effect for *GRK1* disruption, possibly because *GRK1* knockout acts indirectly through SAG by reducing its affinity for the mutant rhodopsin. The nature of this toxic photoproduct is unclear, although we have previously noted the production of a long-lived photointermediate on illumination of both T4K and T17M rhodopsins *in vitro* (Tam et al., 2014). We have also shown that introduction of a disulfide bond increases rhodopsin stability and prevents RD. Interaction with GNAT1, SAG, or GRK1 may similarly stabilize the mutant rhodopsin against adopting abnormal conformations. Notably, GNAT1 and SAG both promote the conversion of metarhodopsin I and metarhodopsin III to metarhodopsin II (Emeis et al., 1982; Schleicher et al., 1989; Zimmermann et al., 2004; Sommer and Farrens, 2006; Frederiksen et al., 2016), and thus these proteins could catalyze the removal of a toxic rhodopsin conformation by promoting the metarhodopsin II conformation.

The requirements for lengthy dark adaptation and rapid light onset suggest that in order to achieve toxic levels, large amounts of the T4K photoproduct must be produced synchronously in order to achieve a threshold level of toxicity; potentially RD requires the retina to exceed some level of production that cannot be counteracted by the GNAT1 and/or SAG activities. The protective effects of pretreatment with prolonged light exposure or gradual light onset shown in Figure 4 may have a similar explanation, with pretreatment initially generating the photoproduct asynchronously so that it cannot be made in sufficient quantities once rapid cycling is initiated due to the majority of the mutant rod opsin lacking a chromophore. An intriguing possibility is that this RD could be an exaggerated form of the light-exacerbated RD present in *SAG* and *GRK1* knockout mice (Chen et al., 1999b, 1999a). The requirement for multiple cycles suggests that the cellular effects of each cycle are cumulative, while the requirement for at least 20 minutes of light exposure suggests the involvement of a relatively slow process, possibly photoconversion of an initial photoproduct, similar to the photoreversal mechanism proposed for light damage involving intense blue light (Grimm et al., 2001). However, more research would be required on all models to definitively connect these RD mechanisms.

Given the requirement for dark adaption and photoactivation to induce RD, the fragmentation and shedding of ROS as early events (preceeding any inner segment changes) and the mitigating effects of other phototransduction proteins, the initial toxic event almost certainly occurs in the ROS. However, our TEM results also suggest that T4K rhodopsin does not denature or aggregate upon photoactivation, as we did not observe disorganized disks. Dysmorphic disks would be expected if the photoactivated mutant assumes a conformation that is less compact or aggregates extensively, similar to results seen for P23H rhodopsin in Figure 3G and previous publications (Haeri and Knox, 2012; Bogéa et al., 2015).

The mechanism underlying this RD is likely shared with several other *RHO* mutants; we have previously found that the *RHO* mutations N15S and T17M induce RD by a similar light and chromophore-exacerbated mechanism in *X. laevis* (Tam and Moritz, 2009; Tam et al., 2014). Like *RHO* T4K, these mutations alter rhodopsin glycosylation and are similarly located in the “chromophore plug” region of the protein. It is also likely that *RHO* T4R dogs and *RHO* T17M mice have similar mechanisms of light-exacerbated disease (Zhu et al., 2004; White et al., 2007). Other light-exacerbated RDs caused by *RHO* mutations have been reported (Budzynski et al., 2010). It remains to be determined whether this chromophore-exacerbated RD mechanism could be responsible for sector RP associated with rhodopin mutantions outside the consensus glycosylation sites (Budzynski et al., 2010).

We suggest caution is warranted in the development of RP treatments. Individuals with sector RP associated with *RHO* mutations should be considered to have at least two possible underlying disease mechanisms that may respond quite differently to pharmacological treatments, as we previously observed for valproic acid (Vent-Schmidt et al., 2017); genotype should be considered when enrolling clinical trial participants. Patients with chromophore-exacerbated disease are unlikely to benefit from pharmacological chaperone treatments, and could potentially benefit from inhibitors of vitamin A metabolism, such as those trialed for Stargardt disease related to mutations in *ABCA4* (Issa et al., 2015; Huang et al., 2021; Kubota et al., 2022). Our results suggest that patients with a T4K-like disease should avoid sudden transitions from darkness to intense light, especially when the light exposure has a duration of minutes or longer and dark adaptation is lengthy. Importantly, such conditions may occur in a standard ophthalmic exam via ophthalmoscopy or photography, and the retinas of *RHO* T4R dogs are documented to be injured by examination with photography procedures involving 60 seconds of light exposure (Cideciyan et al., 2005). In such patients, the brief light exposures possible with modern cameras might be preferable. Other drastic changes in lighting, such as exiting a darkened room into bright daylight, should potentially be avoided. Lifestyle adjustments such as sleeping with curtains open, or use of a sunrise simulator alarm clock, could minimize exposure to sudden changes in light intensity.

## Acknowledgements

The authors wish to thank RS Molday for the gift of mAb2B2, E Weiss for recommending and providing a sample of mAbG8 anti-GRK1, WC Smith for gifts of mAbB630N, anti-GNAT1 polyclonal, and mAb Xarr1-6 anti-SAG, and K Joung for the gift of pDR274. This research was funded by the Canadian Institutes of Health Research (PJT-155937 and PJT-156072) and the National Science and Engineering Research Council (RGPIN-2020-05193).

